# A CURE for synthetic regulation of gene expression: Rapid screening of guide RNA efficacy as a framework for enabling undergraduate research in plant synthetic biology

**DOI:** 10.64898/2026.03.31.715601

**Authors:** Tawni Bull, Liz Carlsen, Noah Hoglund, Josh Blarr, Makenzie Ciernia, Hannah Daughtrey, Keana Gulnac, Zane Kathan, Ben Labovitz, Ryan Lonergan, Mira McDermott, Andrew Medina, Zoey Mikol, Zeke Miller, Kyndal Prahl, Chloe Rifai, Emma Schrems, Futa Shinkawa, Jessie Summerfield, Ethan Thevarajah, Skylar Wagner, Trenton Zimmerman, Arjun Khakhar

## Abstract

Course-based Undergraduate Research Experiences (CUREs) have emerged as a transformative approach to science education, expanding access to authentic research opportunities beyond the traditional undergraduate research assistant (URA) training. By embedding research into a curriculum, CUREs engage a broad and diverse population of students in a classroom environment that emphasizes experimental design, data analysis, and scientific communication. However, this has been difficult to develop for fields such as plant synthetic biology due to the long timescales of plant transformation. One avenue around this problem is to utilize a recent innovation that enables high throughput and rapid screening of gRNA efficacy by leveraging viral-based delivery of guide RNAs (gRNAs). In this work, we develop and validate a CURE with undergraduate students at Colorado State University (CSU). Students worked in teams to design and test efficacy of gRNAs targeting a Cas9-based transcriptional repressor to different regions of the promoters of the three *GIBBERELLIN INSENSITIVE 1* genes (*GID1a, GID1b*, and *GID1c*) in *Arabidopsis thaliana*. Over the semester, students generated and analyzed gene expression data to understand the efficiency of twelve new gRNAs. We further validated CURE student-identified gRNAs with an undergraduate research assistant (URA) that assessed target gene expression and phenotypic outcomes in stable transgenic lines expressing SynTF constructs with the strongest gRNAs from the class. We further describe the curriculum structure to facilitate adoption at other institutions and present student-generated datasets demonstrating the utility of ViN-based screening for identifying effective SynTF gRNAs for plant functional genomics and engineering.

## 1. Introduction

Many studies have shown that course-based undergraduate research experiences (CUREs) are a powerful means to improve students’ engagement with and understanding of Science, Technology, Engineering, and Math (STEM) concepts (1–4). Specifically, CUREs that are embedded into upper-division STEM courses enable students to gain authentic research experiences that may otherwise not be available. However, the experimental realities of certain fields of bioengineering make them challenging to implement as CUREs. This is especially true for plant synthetic biology where the long timescales associated with the generation of transgenic plants often puts such bioengineering outside the scope of a semester long course.

One avenue around this issue is to leverage recent innovations that enable rapid virus-based prototyping of synthetic transcription factor (SynTF)-based gene expression modulation. This system, called Viparinama (ViN), utilizes a positive, single strand plant RNA virus, tobacco rattle virus (TRV), to systemically deliver a truncated guide RNA (gRNA), programming Cas9 to bind and not cut (5), to a transgenic plant line that is pre-engineered to express a host of SynTF components (6). These synthetic components include a constitutively expressed nuclease active Cas9 and the RNA binding protein, PCP, fused to a truncation of the TOPLESS repressor (PCP-TPLN300) (6, 7). TRV-based delivery of the gRNAs, which contains a 3’ modification aiding in the recruitment of the PCP-TPL repressor, into the transgenic lines enables complete assembly of a Cas9-based SynTF at a gene of interest. This allows for modulation of gene expression across plant tissues in a matter of weeks, which contrasts with the months-to-years that it would take to achieve a similar outcome via traditional plant transformation approaches. Such rapid and scalable screening of gRNAs is essential, as SynTFs are known to have variable regulation efficacy based on the binding location of the gRNA target sequence in the promoter (8, 9). Crucially, this approach allows for the study of gRNA efficacy for the regulation of endogenous gene expression in the target tissue, which is not possible with alternative transient approaches such as protoplast transfection. Prior work has validated the utility of ViN for rapidly prototyping SynTF-based expression modulation for functional genetics and plant bioengineering (6).

In this work, we outline a CURE that we designed and validated with a cohort of 19 undergraduate students at Colorado State University (CSU) using ViN as a platform to introduce core concepts in plant synthetic biology. Teams of students, many with no prior lab experience, used ViN to rapidly prototype gRNAs designed to recruit repressors across different regions of the promoters of the three *GIBBERELLIN INSENSITIVE 1* (*GID1a, GID1b*, and *GID1c*) in *Arabidopsis thaliana*. The goal was to identify gRNAs that implement efficient repression of gene expression. From prior work we would expect repression of these genes to generate a dwarfing phenotype (6, 7, 10– 12), providing a tractable and informative readout.

The CURE students received training in paired lab and lecture sections. The lectures focused on the underlying molecular biology, plant developmental biology, viral engineering, and synthetic biology used during the lab section to build a robust theoretical framework necessary to design the experiments and interpret their results. Here, we describe the curriculum of the CURE to facilitate its easy recreation at other institutions. We also describe the data collected by the CURE students to highlight the utility of combining undergraduate education with rapid identification of optimal SynTF gRNAs for functional genetics and plant engineering. Finally, to validate the efficacy of the gRNAs tested by the students, we characterize the expression of the target genes and the resulting phenotypes in transgenic lines stably expressing the SynTFs and an array of the strongest gRNAs identified. The latter work was carried out by a traditionally mentored undergraduate researcher assistant (URA), enabling us to compare the relative strengths of these two approaches to integrating undergraduate training into the research enterprise.

## 2. Materials and methods

### 2.1 CURE outline, materials, and methods Intended students and prerequisite knowledge

This CURE was designed for undergraduate students (sophomores through seniors) that are enrolled in biology-focused programs. This course convers concepts from molecular and cellular biology, genetics, plant biology, virology, and synthetic biology. However, the lecture materials were designed to make it accessible to students with a minimal pre-requisite of a general biology course. While not required, as topics pertaining to the targeted gene family are discussed in detail in the lab and lecture portion, previous or concurrent enrollment in courses that dive more deeply into the underlying biology of the genes being targeted for modulation with SynTFs would likely enhance the students’ experience.

#### Learning Objectives

The learning objectives for this CURE and their respective assessments are outlined in **Supplemental Table S1**. The CURE was designed to introduce and develop students’ understanding of plant synthetic biology and viral engineering, wet lab skills, data analysis, analytical reasoning, and science communication. The following sections describe the CURE lab curriculum. In addition, the lecture materials are included in the supplemental information, although the specifics covered will change based on the gene family being targeted in future iterations of this course.

#### Course Outline: Design, Build, Test, Learn

The CURE was developed as a semester long course (16 weeks) during the Fall semester of 2024. The lecture section met once a week for 50 minutes and met a total of 14 times over the course of the semester. The lab section met twice a week for 1 hour and 50 minutes and met a total of 28 times over the course of the semester. This course followed the structure of the design, build, test, learn cycle common to synthetic biology research (**Fig. 1**). The semester concluded with teams of students practicing their science communication skills via an oral presentation of their workflow, data, and conclusions.

**Figure 1.**
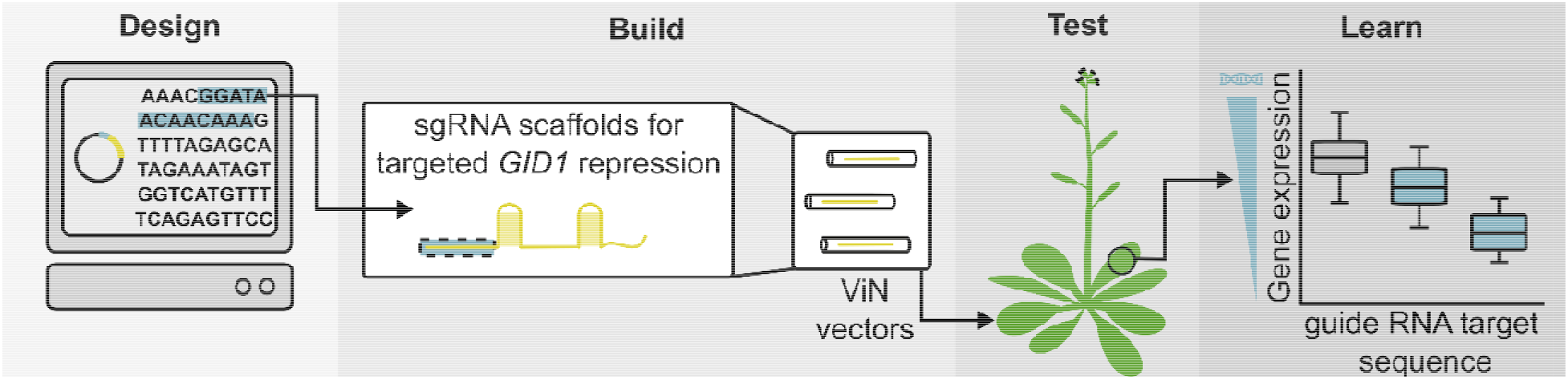
Course outline schematic. A brief summary graphic depicting the structure of the CURE, which follows the classical design, build, test, learn cycle of synthetic biology research.

#### Design: *In silico* design of viral vectors (week 1)

The CURE used a previously published technology, called ViN, that utilizes TRV to deliver gRNA sequences into a plant that stably expresses SynTF components including a nuclease active Cas9 and the RNA binding protein, PCP, fused to the transcriptional repressor, TPLN300 (6). The students were divided into teams of two or three and each team was assigned two gRNA sequences that were pre-designed using CHOPCHOP v3 (13). Each gRNA sequence contained a unique target site and a 3’ recruitment motif (PP7), that were designed to assemble a repressor at the promoter of one of the three *GID1* genes in *Arabidopsis thaliana* (*GID1A, GID1B*, or *GID1c*). Students were given their predetermined gRNA sequences and a previously published TRV2 vector (6) as a template. After one demonstration of how to execute *in silico* design using Benchling (https://www.benchling.com/), they were tasked with designing primers to PCR-amplify DNA fragments capable of being assembled into a TRV2 vector carrying their unique gRNA sequence via Golden Gate assembly. Forward primers were designed to anneal to the *Streptococcus pyogenes* sgRNA handle of the provided TRV2 template DNA (P102) and to add a PaqCI recognition site and a unique *GID1* gRNA sequence to the 5’ end of the amplified DNA fragment. Reverse primers were designed to anneal to the *Flowering Locus T* (*FT*) fragment of the provided TRV2 template DNA (P102) and to add a PaqCI recognition site to the 3’ end of the amplified DNA fragment. The pre-designed gRNA target sequences are listed in **Supplemental Table S2**, primer sequences that the class generated are listed in **Supplemental Table S3**, and the viral vectors and template plasmid are listed with links to their annotated plasmid maps in **Supplemental Table S4**.

#### Build: Golden Gate assembly of viral vectors (weeks 2 through 5)

After designing their viral vectors in Benchling, the students spent the following three weeks focused on building the viral vectors via Golden Gate assembly (14, 15). Specifically, they used their designed primers to PCR amplify DNA fragments using the NEB Q5® High Fidelity DNA Polymerase Master Mix. The amplified product included a unique gRNA sequence, the *S. pyogenes* gRNA handle, the 1xPP7 recruitment motif, and an *FT* fragment all flanked by PaqCI recognition sites. Amplified products were analyzed using gel electrophoresis and purified using the Qiagen Qiaquick Gel Extraction Kit. The students then performed a Level 2 Golden Gate assembly using the Type IIS restriction enzyme PaqCI (15). The DNA components included in the assembly consisted of a TRV2 entry vector (P538) and the insert of the amplified DNA fragment with flanking PaqCI recognition sites to generate compatible 4-bp overhang sequences upon digestion. The assembly was carried out in a one-pot reaction containing a 3:1 insert to vector ratio, the restriction enzyme and T4 DNA ligase. The completed assembly reaction was treated with Plasmid-Safe™ ATP-Dependent DNase (Biosearch Technologies) to remove linear DNA. The assemblies were subsequently transformed into NEB® 10-beta chemically competent *Escherichia coli*, and screened for positive colonies using colony PCR. Colony PCR was performed using a pre-designed set of primers universal to all of the viral vectors being assembled. Positive colonies were grown overnight, and plasmid DNA was isolated using the Qiagen QIAprep Spin Miniprep Kit. For each assembly, purified plasmid DNA from two isolated colonies was sent for Sanger sequencing. The students analyzed their sequencing results using the multiple sequence alignment tool in Benchling. Sequence verified plasmids were transformed into *Agrobacterium tumefaciens* strain GV3101 and used for further downstream analysis to test the efficiency of the gRNAs in plants.

#### Test: Viral-based screening of gRNA target sequences (weeks 6 through 12)

The CURE students tested the efficacy of their unique gRNA sequences at repressing expression of one of the three *GID1* genes via viral delivery into *A. thaliana*. Viral delivery was implemented using the previously described protocol (6). Here, students conducted *Agrobacterium*-mediated co-infiltration of TRV1 and their unique TRV2 vector into mature leaves of *A. thaliana* transgenic plants described above. Seeds of the transgenic line used during the CURE are available upon request. Three plants per gRNA target sequence were infiltrated and one plant per team was co-infiltrated with TRV1 and a no gRNA TRV2 as a control. Plants were grown on a grow rack in the classroom, under fluorescent light set to long day conditions (16hr light/8hr dark). After two weeks, students collected rosette leaf tissue from leaves above the infiltration sites and flash froze them in liquid nitrogen. During the following two classes, students extracted RNA from these tissue samples using the Qiagen RNeasy Plant Mini Kit. The student’s treated the RNA samples with the Invitrogen Turbo DNA-*free*™ Kit and synthesized cDNA using the LunaScript® RT SuperMix Kit. The students measured the relative concentrations of the housekeeping gene, *PP2A*, and the targets genes, *GID1a, GID1b*, and *GID1c*, by performing RT-qPCR using the Luna® Universal qPCR Master Mix. Two technical replicates were performed for each cDNA sample. Primers used for qPCR are listed in **Supplemental Table S5**.

#### Learn: Data analysis and presentation (weeks 13 through 16)

During the two-week period of plant growth, students learned the conceptual basis of qPCR and how the resulting data is analyzed to measure gene expression levels. After collecting the qPCR data, students calculated the relative *GID1* expression levels of each of their samples by normalizing to the housekeeping control gene, *PP2A*. The measurements of *GID1* expression were then compared to the expression in plants treated with the no gRNA control TRV vectors. Students were given two weeks for data analysis and were instructed on various data interpretation strategies implemented in python using Jupyter Lab including statistical tests such as the t-test, ANOVA, and correlation analysis. During the last week of the lab course, teams of students presented their findings and conclusions about their gRNA efficacy based on the *GID1* expression data they collected. They also described areas of possible improvement or pitfalls in their experiments.

### 2.2 URA materials and methods

#### Plasmid construction

The plasmid constructed in this work was built using Golden Gate Assembly and Modular Cloning (14, 15). A Level 1 assembly (15) was performed to build a gRNA array that consisted of the *GID1* gRNA target sequence from each of the three *GID1* genes that resulted in the strongest repression from the class. This was then assembled into a Level 2 (15) T-DNA vector that also contained the SynTF components and transformed into *Agrobacterium tumefaciens* strain GV3101. Both the Level 1 and Level 2 plasmids are listed in **Supplemental Table S4** with a link to its annotated plasmid map.

#### Transgenic line generation

The *A. thaliana GID1* repression transgenic lines described in the work were generated by introducing the SynTF components and best gRNAs targeting the promoters of the three *GID1* genes into the genome of Col-0 via floral dip (16). Seeds from T0 plants were surface sterilized in a solution containing 70% Ethanol and 0.05% Triton-X-100 for 20 minutes. Seeds were immediately rinsed with 95% Ethanol. Seeds were plated on 1/2x MS containing 0.8% PhytoAgar™ (plantmedia Cat. No. 40100072-1) and Kanamycin (50mg/L). Positively selected seedlings were transplanted into soil and grown to maturity and seeds were harvested. Multiple independent T1 lines were used for further experiments. The no gRNA control transgenic lines were used previously (7).

#### Root and hypocotyl phenotyping

Seeds of T1 independent transgenic lines were surface sterilized as described above, plated on 1/2x MS containing 0.8% PhytoAgar™ (plantmedia Cat. No. 40100072-1) and Kanamycin (50mg/L), and stratified in the dark at 4□C for three days. For root assays, plates were placed vertically under short day conditions (9hr light/15hr dark) at 22□C in a Percival growth chamber for eight days. Plates were scanned using an Epson scanner and root length was measured using ImageJ. For hypocotyl assays, plated seeds were light shocked for six hours in a Percival growth chamber set to 22□C and short-day conditions (9hr light/ 15hr dark) and then were wrapped in foil and placed vertically in the correct growth chamber to promote etiolation. After eight days, plates were scanned using an Epson scanner and hypocotyl length was measured using ImageJ.

#### Gene expression analysis

Whole eight-day-old seedlings from the root phenotyping assays of the no gRNA control and new gRNA populations were collected and flash froze in liquid nitrogen. RNA was extracted using the QIAGEN RNeasy Plant Mini Kit (Cat No. 74904). The RNA samples were treated with the TURBO DNA-*Free*™ kit (Invitrogen Cat No. AM1907) to remove genomic DNA. Complementary DNA (cDNA) was synthesized using the LunaScript® RT primer free MasterMix Kit and gene specific primers (**Supplemental Table S5**). Proper removal of the genomic DNA was confirmed with PCR using GoTaq® Green Master Mix (Promega Cat No. M7122) and primers spanning the first and second intron of *GID1a* (**Supplemental Table S5**). Concentrations of *GID1a, GID1b, GID1c*, and *PP2A* were quantified using RT-qPCR performed with the Luna® Universal qPCR Master Mix on the BioRad CFX Opus qPCR machine. Cycle threshold values for *GID1* were normalized to the housekeeping gene, *PP2A*. Two technical replicates were performed for all samples. The gene specific, PCR and RT-qPCR primers used are listed in **Supplemental Table S5**.

#### Data analysis

All data analysis was performed in python using Jupyter Lab (version 3.4.4). The *p*-values reported were calculated using either an ANOVA followed by a Tukey HSD test or Welch’s *t*-test for pairwise comparisons, where appropriate using the scipy package in python. All the data that is presented was plotted using the seaborn package and functions from the matplotlib library (17). All the raw data and code used to analyze the data is available on the following github repository (https://github.com/arjunkhakhar/ViN_CURE_Manuscript).

## 3. Results

### CURE outcomes: Screening gRNAs targeting *GID1* using ViN

In the fall of 2024, students (n = 19) enrolled in the Viral Engineering Laboratory course participated in the CURE described above. Each student was assigned to one of nine teams. Over the first five weeks the students were given the task of designing and building two TRV-based viral vectors, where each carried one gRNA targeted to the promoter of one of the three *GID1* genes in *A. thaliana* (*GID1a, GID1b*, or *GID1c*, **Fig. 2A**). Of the 18 assemblies performed, 15 of them (83%) resulted in isolated bacterial colonies to screen. Of these 15 assemblies, 14 were sequence verified; therefore, the students demonstrated a 78% success rate using Golden Gate modular cloning, which resulted in 14 gRNAs (2 previously tested gRNAs as controls and 12 new gRNAs) to screen *in planta*. During weeks 6 through 8, students were guided through how to deliver their gRNAs into *A. thaliana* plants stably expressing a nuclease active Cas9 and the PCP-TPLN300 transcriptional repressor fusion protein through viral delivery via *Agrobacterium*-mediated co-infiltrations of TRV1 and a TRV2 plasmid encoding a single gRNA. All groups successfully delivered their viral vectors and grew their plants for two weeks until tissue samples were collected and gene expression was measured.

**Figure 2.**
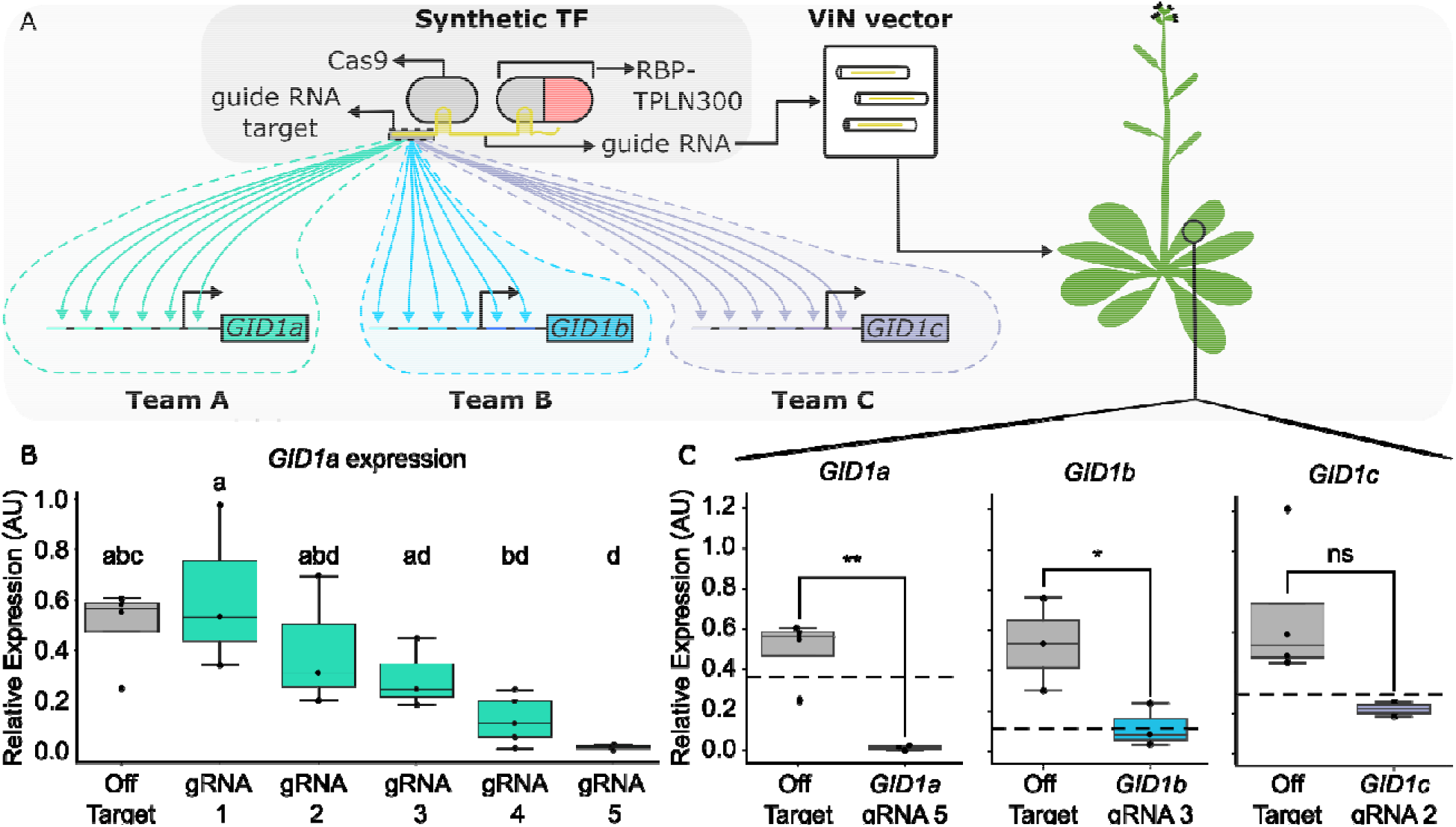
Screening guide RNA efficacy with a CURE. A) A graphic describing the synthetic transcription factor design and the relative positioning of gRNAs across the promoters of *GID1a* (teal), *GID1b* (blue), and *GID1c* (purple). The gRNAs positioned after the transcriptional start site (black arrow) were located in the 5’ UTR of each of the genes. B) Boxplots showing the range of regulation strengths of the gRNAs that were targeted to the promoter of *GID1a*. Different letters represent statistically significant differences (One-way ANOVA followed by Tukey HSD test, *p* < 0.05). C). Boxplots representing the regulation strength for the strongest gRNA for each *GID1a* (teal), *GID1b* (blue), and *GID1c* (purple). Statistical significance calculated using Welch’s two sample *t*-test (*p* < 0.05), * corresponds to *p* < 0.05, ** corresponds to *p* < 0.01, and *** corresponds to *p* < 0.001

During weeks 9 through 12, students measured the relative levels of *GID1* expression for each infiltrated plant. A total of 72 RNA extractions were completed which led to a total of 304 RT-qPCR reactions. Of these RT-qPCR reactions, 244 (80.3%) of them resulted in cycle threshold values for house-keeping genes above background, indicating a successful extraction. Through these reactions, students were able to calculate the *GID1* relative expression levels compared to the housekeeping control, *PP2A*, of their samples (**Fig. 2B, C, Fig. S1A, S1B**). During weeks 13 through 16, students analyzed and visualized their relative expression data using python. This resulted in characterization of the relative regulation implemented by five *GID1a* gRNAs (four new gRNAs and one previously used control, **Fig. 2B**), four new *GID1b* gRNAs (**Fig. S1A**), and five *GID1c* gRNAs (four new gRNAs and one previously used control, **Fig. S1B**). This analysis revealed gRNAs that span a broad range of regulation strengths for all three of the *GID1* genes targeted. This included a percent repression compared to the no gRNA controls that ranged from 6.15% to 97.5% for *GID1a*, 63.8% to 84.7% for *GID1b*, and 13.8% to 68.0% for *GID1c*. Crucially, this screen identified gRNAs that resulted in both stronger and weaker repression than the three *GID1* gRNAs used in previous work (6, 7), which enables future titration of this gene family for functional genetics or plant engineering (**Fig. 2B, S1A, S1B**). While most gRNAs generated weaker or equivalent regulation as previously used gRNAs, the screen identified one *GID1a* gRNA with significantly higher regulation strength (97.5% repression compared to the no gRNA control) compared to the gRNA previously used (32% repression compared to the no gRNA control, **Fig. 2C**). This data highlights the utility of using a CURE for rapid prototyping of the relative efficacy of gRNAs for SynTF based expression modulation *in planta*.

### Undergraduate research assistant (URA) outcomes: Characterizing novel gRNAs targeting *GID1* in stable transgenic lines

To further investigate the power of ViN-based screening of gRNA sequences through a CURE, we built and characterized *A. thaliana* lines that stably expressed Cas9, PCP-TPLN300, and a gRNA array that consisted of the three strongest guides identified by the CURE students (**Fig. 3A**). This work was largely conducted by a single undergraduate researcher that was more traditionally mentored and trained in our lab group and took approximately one year to complete. To validate the efficacy of the improved gRNAs identified by the CURE students, we measured the expression levels of *GID1a, GID1b*, and *GID1c* across multiple independent transformation events. We observed a significant decrease in expression strength of each gene when compared to the no gRNA population (**Fig. 3C-E**). In addition, when we compared the repression strength of the new gRNAs to the old guides (7), we observed a significant increase in repression strength for *GID1a* (24% stronger repression, *p* < 0.001, **Fig. 3C**). We also observed stronger repression for *GID1b* (16% stronger repression, **Fig. 3D**) and slightly weaker repression for *GID1c* (2% weaker expression, **Fig. 3E**), although neither was significant. While each new guide resulted in stronger or equivalent regulation during the CURE, the increase in regulation strength for both *GID1b* and *GID1c* was minimal across stable lines.

**Figure 3.**
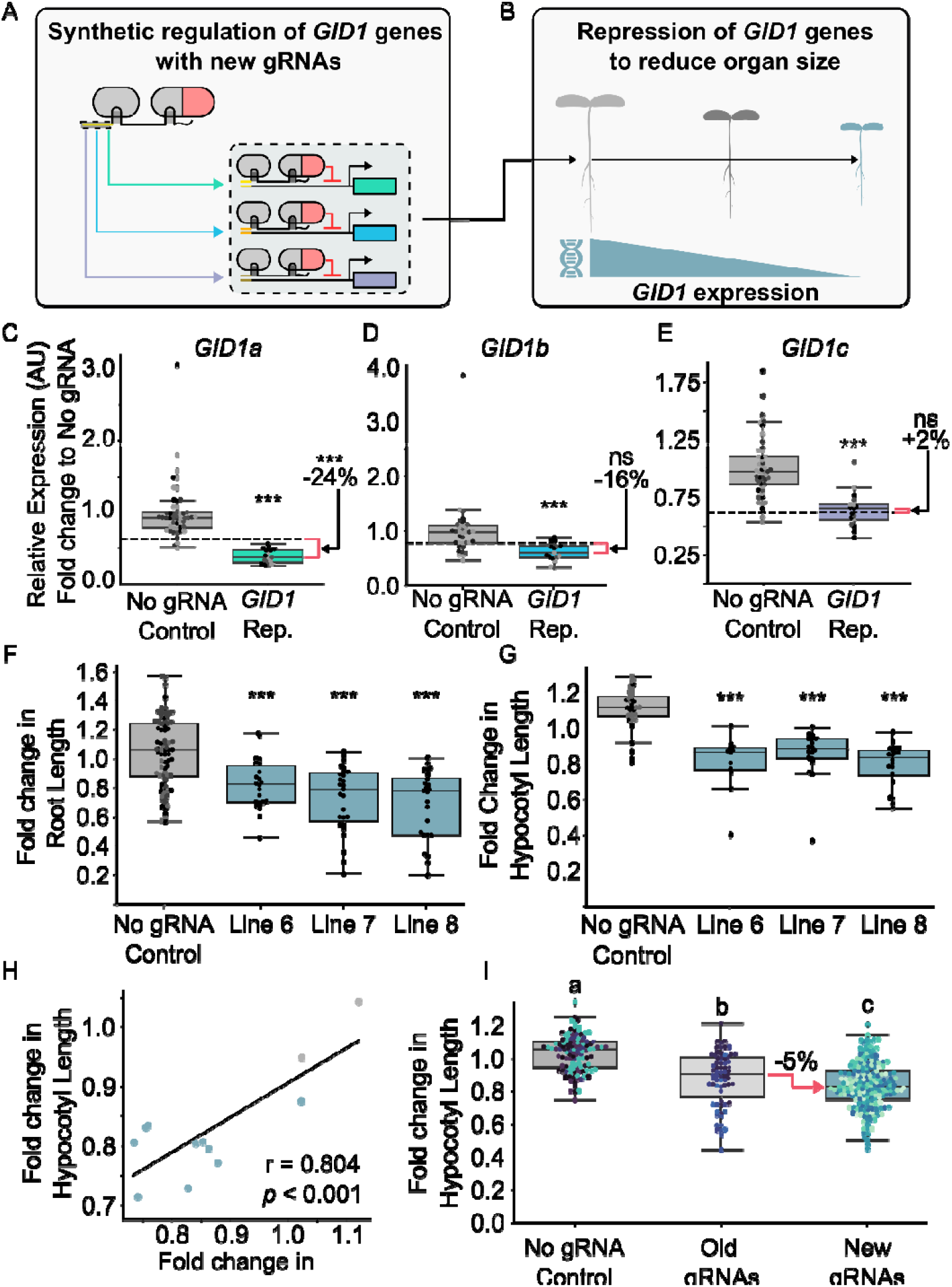
Validating gRNA efficacy in stable transgenic lines. A) Schematic of the genetic circuit designed to implement SynTF-based repression on the three *GID1* genes in *A. thaliana*. B). Schematic depicting the expected engineered phenotype generated with the new gRNAs (blue). C-E) Relative expression levels of *GID1a* (C, n=3), *GID1b* (D, n=3), and *GID1c* (E, n=3) compared to a population of no gRNA controls (gray, n=4). The blacked dashed line represents the average expression level of each gene across a population of plants stably expressing the old gRNA targets sequences (7). F, G) Boxplots representing the fold change in root length (F) or hypocotyl length (G) compared to the no gRNA control population (gray) from three independent transgenic events stably expressing the new gRNAs. H) Pearson’s correlation representing the relationship between the fold change of hypocotyl and root lengths compared to the no gRNA control population. I) Boxplots representing the fold change in hypocotyl length compared to the no gRNA control population (light gray, n=6) of the transgenic populations stably expressing the old gRNAs (dark gray, n=5), and the new gRNAs (blue, n=10). The red arrow represents the percent reduction in size between the old (dark gray) and new (blue) gRNAs. Across all boxplots, each dot of the same color represents an independent biological replicate. Statistical significance calculated using Welch’s two sample *t*-test (*p* < 0.05, * corresponds to *p* < 0.05, ** corresponds to *p* < 0.01, and *** corresponds to *p* < 0.001) or a one-way ANOVA followed by Tukey HSD test, *p* < 0.05.

Previous work has demonstrated that targeting each of the three *GID1* genes in *A. thaliana* with a SynTF repressor results in a decrease in organ size across the plant body (7). Therefore, to investigate the relative efficacy of the improved gRNAs at producing expected phenotypes, root and hypocotyl lengths were measured for ten independent transgenic events (**Fig. 3F, 3G**). As controls, we used the previously reported no gRNA control lines that retain the SynTF components but lack the gRNAs (7). We observed significant decreases in size of both tissues across multiple independent lines that were phenotyped (**Fig. S2A, S2B, S2C, S2D**). The *GID1* repression lines had reductions in root length that ranged from 2% to 36% and reduction in hypocotyl lengths that ranged from 12.5% to 28.6% when compared to the no guide RNA population. Additionally, we observed a strong correlation between the fold change in hypocotyl length and the fold change in root length, demonstrating consistent dwarfing across tissues (**Fig. 3H**), as reported previously (7).

To investigate whether the new gRNAs identified by the CURE students resulted in stronger phenotypes, we compared the fold change in organ size across hypocotyl tissue to previously reported data implementing the same SynTF strategy for repression of *GID1s* (7). Stably expressing the improved gRNAs from the CURE resulted in hypocotyls with a larger reduction in organ size across the population when compared to the gRNAs characterized previously (5% decrease in hypocotyl length across lines, *p* = 0.0017, **Fig. 3I**). The new gRNA population also shows a more consistent dwarfing effect across the whole population (CV = 5.4%) when compared to the old gRNA population (CV = 15.2%, **Fig. S3A**). Additionally, the variance of hypocotyl length within each independent event is similar across all genotypes (**Fig. S3B**). Therefore, the difference in CV between the new and old gRNA populations is driven by variation in hypocotyl length within the genotype. Taken together, these results validate the utility of CURE-based screening of gRNAs to enable rapid identification of gRNAs for functional genetics and plant engineering with SynTFs.

## 4. Discussion

One of the biggest challenges to training the next generation of plant synthetic biologists is the lack of interdisciplinary research opportunities that equip students with both the theoretical frameworks and experimental skills necessary to pursue plant bioengineering research in industry or academia. This is in part due to the general lack of courses that cover topics in both bioengineering and plant biology, and in part due to the limited number of labs pursuing such research (18). However, there is an urgent need to expand the number of trainees in this space to generate the agricultural innovations necessary to secure our food systems and bioindustries from the threat posed by climate change. CUREs represent one mechanism that can sustainably fill this gap. However, the nature of experimental timelines associated with plant transformation has often stymied their creation for plant synthetic biology.

The curriculum we present here offers a relatively low investment approach to training students on the experimental fundamentals for plant synthetic biology. Students leave with proficiency in molecular cloning of plasmids, *Agrobacterium*-mediated transient expression, RT-qPCR based expression analysis, and python-based data analysis and visualization. They also gain an understanding of a range of plant synthetic biology topics including SynTFs, plant viral vector engineering, strategies for biocontainment, plant molecular biology, and plant physiology. The team-based learning students engage in creates peer support networks within the group and thereby lowers the burden on the course’s instructors, enabling larger class sizes.

We taught this CURE at CSU for the first time in 2024 and chose to focus on the *GID1* gene family in *A. thaliana*, as we had a set of previously validated gRNAs for these genes which we used as positive controls (6, 7). We show that over the course of the semester the students were able to screen multiple gRNAs for the three different genes and identify gRNAs that implemented robust regulation for each gene. The efficacy of gRNA-directed SynTFs in gene regulation and dwarfing phenotypes was independently validated the following year by a traditionally mentored undergraduate research assistant. This student, who had undergone a year of prior training, took a year to generate this data (**Fig. 4**). This highlights the effectiveness of utilizing ViN vectors over traditional prototyping of gRNA efficacy via generation of stable transgenic lines. It also proves that this CURE was able to generate useful experimental data about gRNA functionality, while also creating valuable research experience for the undergraduate students enrolled in the course.

**Figure 4.**
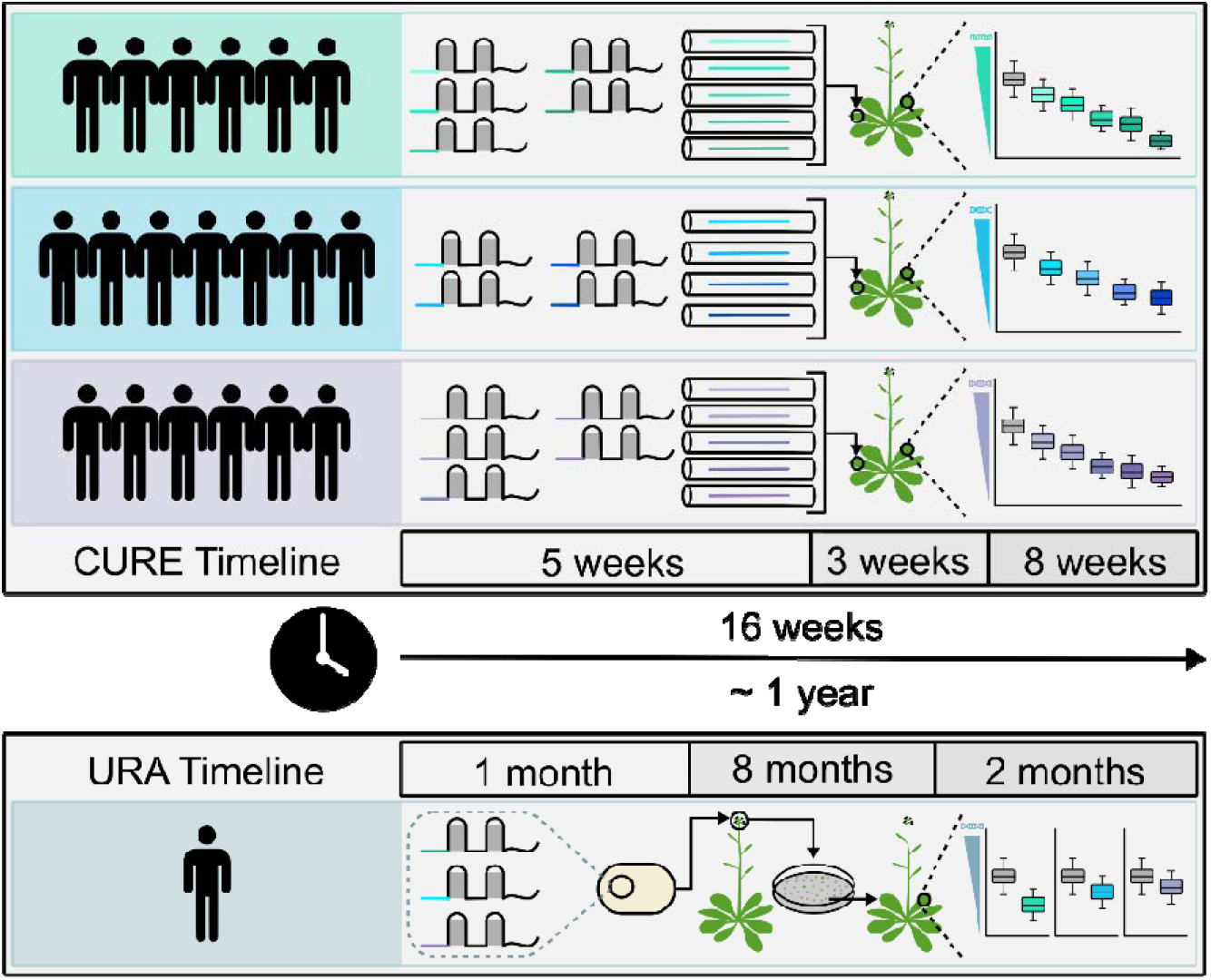
Timeline of screening gRNAs utilizing ViN implemented into a CURE compared to screening gRNAs via a URA generating transgenic lines stably expressing a Cas9-based SynTF and gRNAs.

While the initial instance of the CURE we taught was focused on *GID1* regulation of organ size, the ViN system could be applied to prototype SynTF-based expression modulation of any gene in future iterations of this course by altering the target sites in the gRNAs. Additionally, we have previously shown that this system could be applied to a range of different plant species (6) further expanding the scope of functional genetics questions or bioengineering goals that the CURE could be focused on. We hope to study other genes in the gibberellin signaling pathway in tomato and sorghum in future iterations of this CURE. As such, this CURE represents a flexible way for plant science researchers to provide students with interdisciplinary research opportunities in synthetic biology while also generating useful functional genetics data for their research programs.

## Material Availability

All course materials are available on github (https://github.com/arjunkhakhar/ViN_CURE_Manuscript). All plasmids generated in this work will be available on Addgene. Seeds of transgenic lines used during the CURE are available upon request.

## Data Availability

Supplementary Data is available at SYNBIO online. All data and the code used for data analysis and plotting are available at the following github repository: https://github.com/arjunkhakhar/ViN_CURE_Manuscript

## Author Contribution

T.B. and A.K. designed the CURE. T.B. and N.H. taught the laboratory section. A.K. taught the lecture section. T.B., L.C., N.H., J.B., M.C., H.D., K.G., Z.K., B.L., R.L., M.M., A.M., Z.M., K.P., C.R., E.S., F.S., J.S., E.T., S.W., T.Z. assisted with experiments and analyzed the experimental data. T.B. and A.K. wrote the manuscript.

## Acknowledgments

We would like to thank DeeDee Wright for taking the time to evaluate and give us feedback on the first iteration of this course.

