## Supplemental Information for "A CURE for synthetic regulation of gene expression: Rapid screening of guide RNA efficacy as a framework for enabling undergraduate research in plant synthetic biology"

### Supplemental Figures

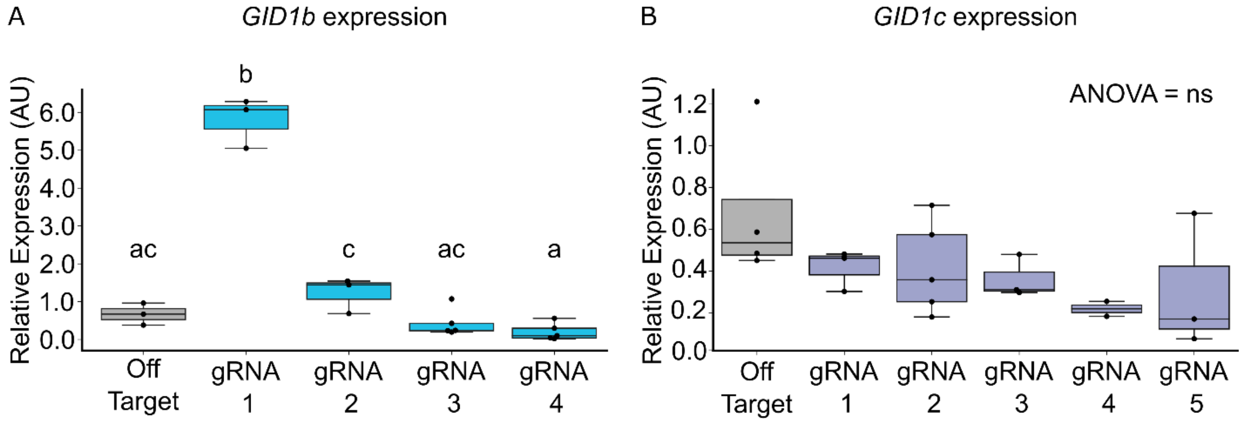


**Supplemental Figure S1. Efficacy of different *GID1* gRNA target sequences.** Boxplots representing the relative expression of either *GID1b* (A) or *GID1c* (C) normalized to a housekeeping gene. The expression levels were generated by targeting different regions of each promoter with unique gRNA sequences that were built and tested using ViN by the CURE students. Different letters represent statistically significant differences (One-way ANOVA followed by Tukey HSD test, $p < 0.05$).

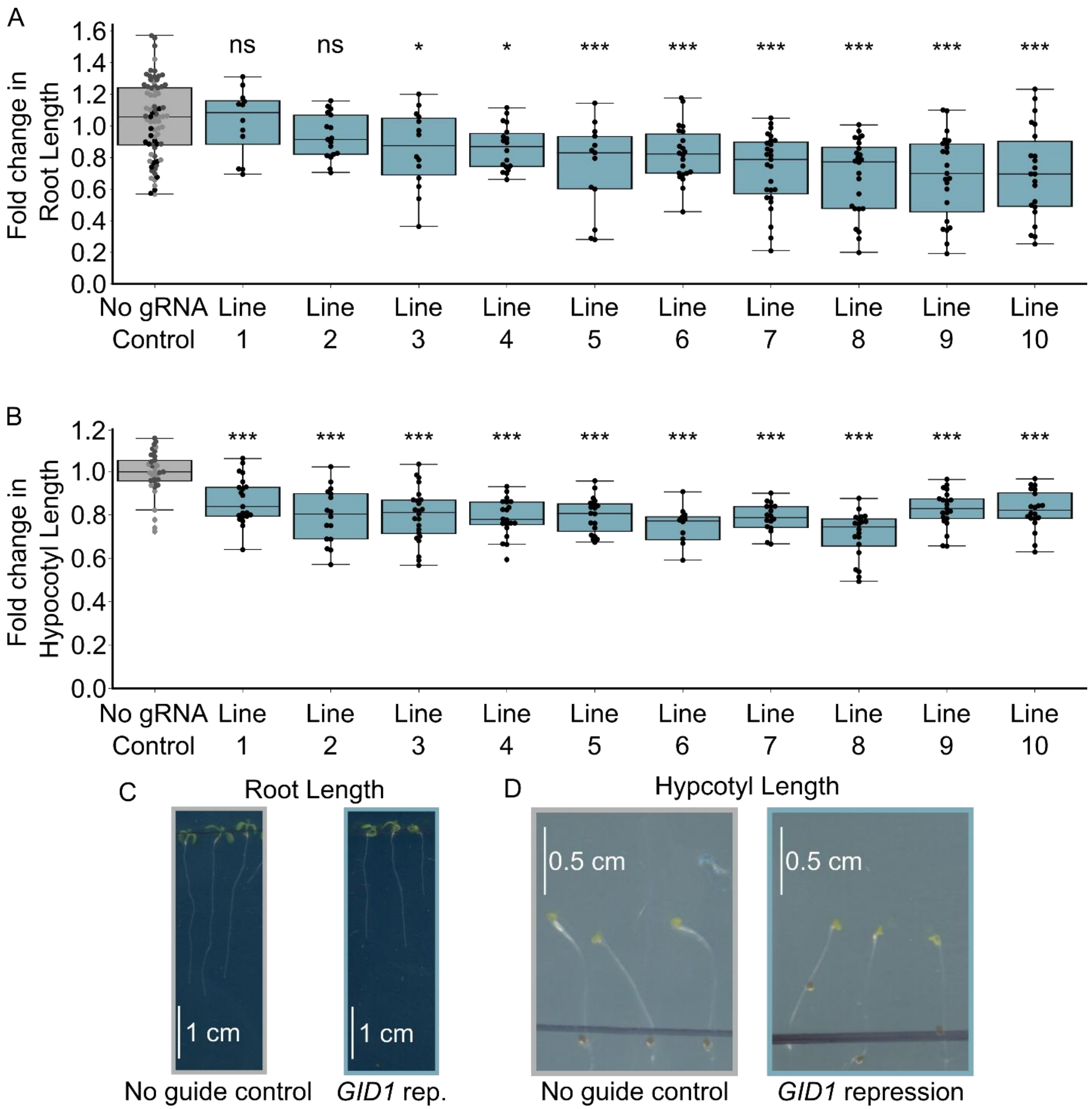


**Supplemental Figure S2. Reduction in organ size of the new gRNA population.** Boxplots representing the fold change in root (A) or hypocotyl (B) length compared to the control population of 10 independent transgenic events expressing the new gRNA array consisting of the strongest gRNAs identified by the CURE. Every dot of the same color within each boxplot corresponds to an independent biological replicate of the same genotype. C) Representative scans of root lengths of the no guide RNA control (left, gray) and the new *GID1* repression line 8 (right, blue). D) Representative scans of hypocotyl lengths of the no guide RNA control (left, gray) and the new *GID1* repression line 8 (right, blue). Significance calculated via a one-way ANOVA followed by a Tukey HSD test comparing each independent line to the control (ns = not significant, *$p < 0.05$, **$p < 0.01$, ***$p < 0.001$).

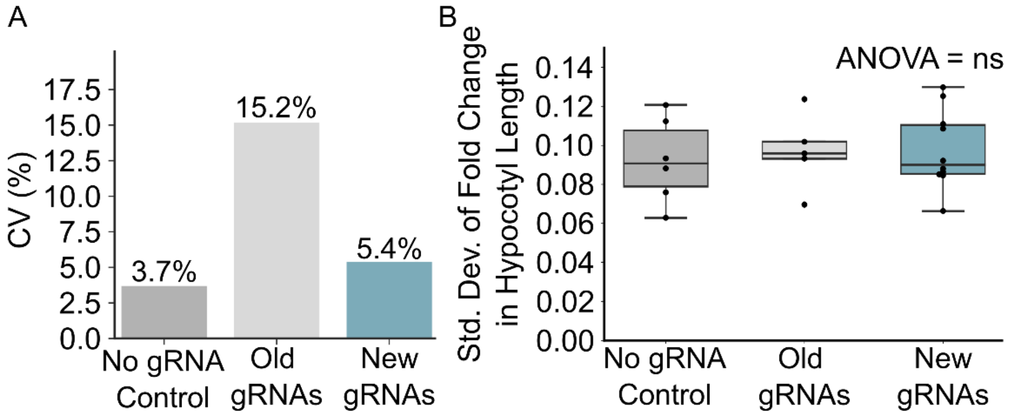


**Supplemental Figure S3. Consistency of the reduction in hypocotyl length across different genotypes.** A) Barplot showing the coefficient of variation (CV) for the no gRNA control population (dark gray), the old gRNA population (light gray), and the new gRNA population (blue). B) Boxplots depicting the standard deviation of the fold change in hypocotyl length to the control population for the no gRNA population (dark gray), the old gRNA population (light gray), and the new gRNA population (blue). Each dot represents a biological replicate for each genotype.

### Supplementary Tables

**Supplemental Table S1.** CURE student learning objectives

| **Student Learning Objectives** |
|---|
| *Lecture Learning Objectives* |
| 1. Describe the mechanisms used by viruses to infect, replicate, and spread in different hosts. |
| 2. Describe the pervasive nature of viruses in all forms of life and of the various strategies they adopt, from commensal to pathogenic. |
| 3. Explain how viruses are used as effective tools to study and engineer biology. |
| 4. Articulate the importance of containment of engineered viral vectors. |
| 5. Explain how plant viral vectors (RNA viruses) are used to study gene expression and edit genomes with RNAi based knockdown. |
| 6. Explain how plant biology can be studied using synthetic transcription factors deployed on viral vectors. |
| 7. Describe how computational techniques are used to domesticate viruses into effective vectors |
| 8. Explain how Adenoviral vectors and Lentiviral vectors can be used to study neural connectivity and to facilitate therapeutic gene delivery and oncolytics. |
| 9. Explain how bacteriophages are used to design antibacterials and how they are used to study protein-protein interaction with phage display. |
| 10. Explain how viruses are used for protein production and how this is used to produce vaccines in plants or virus-like particles for functionalized biomaterials. |
| 11. Specify appropriate strategies to contain viral vectors in both academic and industrial settings. |
| 12. Articulate career opportunities available in viral engineering. |
| *Laboratory Learning Objectives* |
| 1. Use the web-browser based DNA design tool Benchling (https://www.benchling.com/) to design viral vectors and describe the molecular cloning procedure to assemble and deploy them |
| 2. Use Goldengate Assembly and modular cloning to construct viral vectors |
| 3. Use Sanger sequencing to verify vectors via *in-silico* multiple sequence alignment in Benchling (https://www.benchling.com/) |
| 4. Use Agrobacterium to deliver viral vectors to plants |
| 5. Use RNA extraction and qPCR kits to test gene expression modifications |
| 6. Analyze experimental results related to viral engineering and make conclusions based on controls |

**Supplemental Table S2.** Guide RNA target sequences screened by the CURE students. The bold sequences are the gRNAs that resulted in the strongest regulation.

| Target | Guide RNA | Sequence (5' to 3') |
|---|---|---|
| *GID1a* | 1 | TGTTGGGTCATGGG |
| | 2 | GTCAATTTTATTCG |
| | 3 | AGGGATGAGTAGGG |
| | 4 | TGCGCGCATAAGTT |
| | **5** | **GGATAAACAACAAA** |
| *GID1b* | 1 | AGCACTAATGAGAG |
| | 2 | GAATTAGGAAGCAG |
| | 3 | TACACATGACACCG |
| | **4** | **CTAATAATATTATG** |
| *GID1c* | 1 | AAGAATATCGGCGT |
| | 2 | TGCGAACAGATTAA |
| | 3 | TCAGAATACTAGAA |
| | **4** | **CGACTGTTGTTGGG** |
| | 5 | CACCCTTGGGAGCA |

**Supplemental Table S3.** Primers designed by the CURE student used to amplify Golden Gate compatible fragments containing each gRNA sequence. The underlined sequence represents the PaqCI restriction sequence and four base pair overhang sequences. The bold sequence represents the gRNA target sequence. The remaining sequence annealed to the TRV2 template provided to the class.

| Primer Name | Sequence (5' to 3') |
|---|---|
| PaqCI_AAAC-GID1a-1-fwd | gtcacctgcttttaaac**TGTTGGGTCATGGG**GTTTTAGAGCTAGAAATAGCAAG |
| PaqCI_AAAC-GID1a-2-fwd | gtcacctgcttttaaac**GTCAATTTTATTCG**GTTTTAGAGCTAGAAATAGCAAG |
| PaqCI_AAAC-GID1a-3-fwd | gtcacctgcttttaaac**AGGGATGAGTAGGG**GTTTTAGAGCTAGAAATAGCAAG |
| PaqCI_AAAC-GID1a-4-fwd | gtcacctgcttttaaac**TGCGCGCATAAGTT**GTTTTAGAGCTAGAAATAGCAAG |
| PaqCI_AAAC-GID1a-5-fwd | gtcacctgcttttaaac**GGATAAACAACAAA**GTTTTAGAGCTAGAAATAGCAAG |
| PaqCI_AAAC-GID1b-1-fwd | gtcacctgcttttaaac**AGCACTAATGAGAG**GTTTTAGAGCTAGAAATAGCAAG |
| PaqCI_AAAC-GID1b-2-fwd | gtcacctgcttttaaac**GAATTAGGAAGCAG**GTTTTAGAGCTAGAAATAGCAAG |
| PaqCI_AAAC-GID1b-3-fwd | gtcacctgcttttaaac**TACACATGACACCG**GTTTTAGAGCTAGAAATAGCAAG |
| PaqCI_AAAC-GID1b-4-fwd | gtcacctgcttttaaac**CTAATAATATTATG**GTTTTAGAGCTAGAAATAGCAAG |
| PaqCI_AAAC-GID1c-1-fwd | gtcacctgcttttaaac**AAGAATATCGGCGT**GTTTTAGAGCTAGAAATAGCAAG |
| PaqCI_AAAC-GID1c-2-fwd | gtcacctgcttttaaac**TGCGAACAGATTAA**GTTTTAGAGCTAGAAATAGCAAG |
| PaqCI_AAAC-GID1c-3-fwd | gtcacctgcttttaaac**TCAGAATACTAGAA**GTTTTAGAGCTAGAAATAGCAAG |
| PaqCI_AAAC-GID1c-4-fwd | gtcacctgcttttaaac**CGACTGTTGTTGGG**GTTTTAGAGCTAGAAATAGCAAG |
| PaqCI_AAAC-GID1c-5-fwd | gtcacctgcttttaaac**CACCCTTGGGAGCA**GTTTTAGAGCTAGAAATAGCAAG |
| PaqCI_CAGT-rv | gtcacctgcttttactgTTGGCCATAAGTAACCTTTAG |

**Supplemental Table S4.** A list of plasmids that were constructed for this work with links to their annotated plasmid maps.

| Plasmid Number | Plasmid Name | Link to Annotated Plasmid Map |
|---|---|---|
| P538 | TRV2 Entry Vector | https://benchling.com/s/seq-lEQWWJojNCRMlBc6wIlI?m=slm-75DzTfVZhUxNjlfFeF1F |
| P182 | TRV1 | https://benchling.com/s/seq-ZCZd2jVlviYBdk01Ybgp?m=slm-sL1w43YmTxrskkmpVZJy |
| P1651 | TRV2_pAtGID1a_gRNA_target_truncated_1 | https://benchling.com/s/seq-ARTaKvCDQy2tCkHi4riL?m=slm-Xmh7Snrc5tTjgK8hEJLH |
| P1655 | TRV2_pAtGID1a_gRNA_target_truncated_2 | https://benchling.com/s/seq-4oqBVVgA0y3q7CYuVkEc?m=slm-oOc2oe8imnzFTIy7xmub |
| P102 | TRV2_pAtGID1a_gRNA_target_truncated_3 | https://benchling.com/s/seq-jUgFxCy6HOC8JE29kpgP?m=slm-Cjjoxeekfcx6L6IlNdzO |
| P1652 | TRV2_pAtGID1a_gRNA_target_truncated_4 | https://benchling.com/s/seq-R4cHPqWbGQuzAVRXk4g9?m=slm-mbp8auUtqU6bcSJyAmaR |
| P1654 | TRV2_pAtGID1a_gRNA_target_truncated_5 | https://benchling.com/s/seq-h5Yw8REPiw8cV4nirkm3?m=slm-cxg6l90YdinJPOl9P1Lg |
| P1660 | TRV2_pAtGID1b_gRNA_target_truncated_1 | https://benchling.com/s/seq-oo5Nz8FoD3jwZTA0KNCm?m=slm-tEFPRbPbmVQ6OV4yh1U3 |
| P1659 | TRV2_pAtGID1b_gRNA_target_truncated_2 | https://benchling.com/s/seq-w3KPaukjxIAVkuBdDTIO?m=slm-ZsSB2Ng5XULCKWZ4Uwg2 |
| P1656 | TRV2_pAtGID1b_gRNA_target_truncated_3 | https://benchling.com/s/seq-bgWgzAYal64m4NT3SqWd?m=slm-b4ENQH80jTfxRFfpSeoL |
| P1657 | TRV2_pAtGID1b_gRNA_target_truncated_4 | https://benchling.com/s/seq-SFkaKh2zuiSBvf05q5Wx?m=slm-n5N3ONAsYCjEwXg8gQC3 |
| P104 | TRV2_pAtGID1c_gRNA_target_truncated_1 | https://benchling.com/s/seq-cM3MKbNqvuBEc7HPx9VJ?m=slm-SLoRDxCmCYhPu3c98JmL |
| P1662 | TRV2_pAtGID1c_gRNA_target_truncated_2 | https://benchling.com/s/seq-uYJK54hsDhArXt5DmzsE?m=slm-2Da5wYUnDCSGdT2ojfVm |
| P1664 | TRV2_pAtGID1c_gRNA_target_truncated_3 | https://benchling.com/s/seq-KJtGk6ZoG1998OJFnRQk?m=slm-uotEzLGCDZEfBzDKdrMJ |

| Plasmid Number | Plasmid Name | Link to Annotated Plasmid Map |
|---|---|---|
| P1661 | TRV2_pAtGID1c_gRNA_target_truncated_4 | https://benchling.com/s/seq-sy917JYNQjrCPaYKo0QE?m=slm-pNJZwR0ppMHXyxypWgGu |
| P1665 | TRV2_pAtGID1c_gRNA_target_truncated_5 | https://benchling.com/s/seq-GwUOeIZmkZQ9aiLpivbT?m=slm-uF7tPPzalCd5nSmmF1cz |
| P1741 | pMOD_D-p35s-GID_guides_3-t35s | https://benchling.com/s/seq-W9uKsfGMAZFsc7F3MfRh?m=slm-psS6t8rcZa8nJfv268yK |
| P1802 | pTRANS_220d - pUBQ10:NLS-Cas9-tHSP - pGmUbi:NLS-MCP-DREB2A-tUBQ1 - p35s:NLS-PCP-TPLN300:tNos - p35s-GID_guides_3-t35s | https://benchling.com/s/seq-4nrXKgXM1BuAfaCG6SsW?m=slm-N7IegWEd5xqoxKIYyjdR |

**Supplemental Table S5.** A list of primers used for gene expression analysis.

| Primer Name | Primer Sequence (5' to 3') | Use |
|---|---|---|
| AtGID1a qPCR f | GGCTCAAGAAAGCGGGTCAAGAG | RT-qPCR |
| AtGID1a qPCR r | GCGTTTACAAACGCCGAAATCTCATCC | RT-qPCR |
| AtGID1b_qPCR-f | ATGTTTGGTGGACAGGAGAG | RT-qPCR |
| AtGID1b_qPCR-r | CGGTAGATAAGCCCTCCAATAC | RT-qPCR |
| AtGID1c_qPCR-f | GTTTGGAGGGACCGAAAGAA | RT-qPCR |
| AtGID1c_qPCR-r | AGGAAGAAACGCTCTCCAATAC | RT-qPCR |
| PP2A_q-f | AACGTGGCCAAAATGATGC | RT-qPCR |
| PP2A_q-r | AACCGCTTGGTCGACTATCG | RT-qPCR |
| GID1a_exon1toexon2_junction-f | CACTTCTCGACTTGCAAATTC | PCR to check gDNA contamination |
| GID1a_exon1toexon2_junction-r | ATTGTAGGCTACTTTGAAGTTGG | PCR to check gDNA contamination |
| AtGID1a-r | CCCAACAGTTGCTTTCTC | Gene specific cDNA synthesis |
| AtGID1b-r | CCATAAGACAATGAAAGTGATCATTG | Gene specific cDNA synthesis |
| AtGID1c-r | GTAGAAGCCAATAGTGGCTTG | Gene specific cDNA synthesis |
